# Lactate Oxidase (LctO) Acts as a Metabolic Checkpoint Restraining *Streptococcus pneumoniae* Invasion of Respiratory Epithelial Barriers

**DOI:** 10.64898/2026.07.15.738724

**Authors:** Ana G. Jop Vidal, Kenichi Takeshita, Landon Murin, Verónica Roxana Flores-Vega, Babek Alibayov, Roberto Rosales-Reyes, Jason W. Rosch, Eva Bengten, Jorge E. Vidal

**Author notes:** Address correspondence to: Jorge E. Vidal, PhD, 2500 North State Street, Jackson Mississippi 39216.

## Abstract

*Streptococcus pneumoniae* rapidly translocates across polarized human bronchial epithelial barriers, with viable bacteria recovered from the basolateral compartment within 1 h post-infection. Disruption of the pyruvate node through combined deletion of pyruvate oxidase (*spxB*) and lactate oxidase (*lctO*) markedly enhanced transmigration of *S. pneumoniae* across polarized Calu-3 monolayers without causing early cytotoxicity or loss of monolayer integrity. This hyper-invasive phenotype was conserved in the TIGR4 and EF3030 background and under air-liquid interface conditions. Importantly, single Δ*lctO* mutants exhibited significantly greater translocation than Δ*spxB* mutants or wild-type strains across bronchial (Calu-3), alveolar (A549), and pharyngeal (Detroit 562) epithelial models. Enhanced translocation correlated with increased bacterial adherence but was independent of capsule expression, extracellular H₂O₂ production, pneumolysin, or tight junction disruption, as evidenced by stable transepithelial electrical resistance (TEER), lack of caspase-3/7 activation, and minimal IL-18 release at early time points. High-resolution confocal microscopy revealed intracellular Δ*lctO* pneumococci localized within N-acetylglucosamine/sialic acid (GN/SA)-containing compartments as early as 1 h post-infection. In murine macrophages, Δ*lctO* mutants were phagocytosed at rates similar to wild-type bacteria but induced greater pneumolysin-dependent cytotoxicity at 24 h. These findings demonstrate that LctO functions as a metabolic checkpoint that restrains pneumococcal invasion of respiratory epithelia, revealing a previously unrecognized role for lactate oxidase in controlling the transition from colonization to invasive disease.

**Importance:** This study identifies lactate oxidase (LctO) as a critical metabolic checkpoint that restrains *Streptococcus pneumoniae* invasion of respiratory epithelial barriers. By linking pyruvate node metabolism to the control of transmigration, these findings reveal a novel mechanism by which central carbon metabolism regulates pneumococcal virulence and the transition from colonization to invasive disease.

## Introduction

*Streptococcus pneumoniae* (the pneumococcus) remains a leading cause of community-acquired pneumonia, otitis media, meningitis, and bacteremia worldwide^1^. A hallmark of pneumococcal disease is the bacterium’s remarkable ability to cross mucosal and epithelial barriers to invade the pulmonary parenchyma, including bronchioles and alveolar spaces^1,2^. In severe cases, pneumococci breach endothelial barriers, gaining access to the bloodstream and resulting in life-threatening bacteremia and disseminated infection^3^. While extracellular colonization of the nasopharynx and upper respiratory tract is relatively well understood, the cellular and molecular mechanisms that enable intracellular survival, translocation, and dissemination of pneumococci remain incompletely defined, largely due to technical limitations of existing *in vitro* models for studying these transient intracellular lifestyles.

Host cell receptors and specific pneumococcal surface components mediate initial attachment and invasion of epithelial surfaces^1,2,4^. Key interactions include the binding of phosphorylcholine (ChoP) residues on the pneumococcal cell wall to the platelet-activating factor receptor (PAFr) and the engagement of pneumococcal surface protein C (PspC/CbpA) with the polymeric immunoglobulin receptor (pIgR). These interactions facilitate adherence, uptake, and transcytosis across nasopharyngeal, respiratory, and endothelial cells^5,6^. Following adhesion, dynamin-dependent endocytosis allows internalization of pneumococci as early as 30 minutes post-inoculation. A subpopulation of internalized bacteria avoids lysosomal degradation by trafficking through recycling endosomes, ultimately enabling basolateral translocation of viable organisms^7^. In contrast, traversal of the blood-brain barrier, modeled using human brain microvascular endothelial cells, involves both dynamin-dependent and –independent endocytic routes^8^.

In addition to receptor-mediated uptake of intact bacteria, a PAFr-independent, macropinocytosis-like pathway has been described for internalization of pneumococcal cell wall fragments (peptidoglycan-teichoic acid). While the PAFr-dependent route elicits strong inflammatory responses, the receptor-independent pathway appears immunologically silent^9^.

Contact between *S. pneumoniae* and bronchial epithelial cells triggers rapid, LytA-dependent capsule shedding^10,11^. Although decapsulation facilitates initial adhesion and entry, it is insufficient for efficient invasion, as non-encapsulated mutants generally show attenuated virulence *in vivo*. We recently reported that pneumococci can rapidly re-encapsulate upon reaching the alveolar compartment, a process partially driven by labile heme released from damaged capillaries—a signal absent in bronchial epithelium^10^. These observations highlight the importance of rapid metabolic adaptations that support virulence. In particular, enzymes of the pyruvate node have been implicated in maintaining NADH/NAD⁺ redox balance and modulating acetyl-CoA availability for capsule biosynthesis^12^. This is consistent with the function of lactate oxidase (LctO), which recycles lactate to pyruvate, thereby supporting H₂O₂ production, a factor linked to pneumococcal virulence^13^.

Despite these advances, the bacterial factors that orchestrate rapid phenotypic switching upon epithelial contact to promote intracellular invasion and basolateral translocation remain poorly defined. Metabolic enzymes involved in lactate utilization play conserved roles in invasive disease across Gram-positive pathogens. In *Streptococcus pyogenes*, LctO oxidizes host-derived lactate to pyruvate, generating H₂O₂ that promotes tissue damage and invasion. Similarly, lactate metabolism supports persistence and systemic spread of *Staphylococcus aureus* under host-imposed stresses such as nitric oxide exposure. Here, we demonstrate that LctO, which irreversibly converts L-lactate to pyruvate in *S. pneumoniae*, functions as a critical metabolic checkpoint that restrains rapid epithelial invasion and transmigration.

## Results

### Rapid translocation of pneumococci across polarized human bronchial cells

We initially determined the optimal differentiation period for human Calu-3 cells by monitoring stable transepithelial resistance (TEER). At 10 days post-seeding, the median TEER (∼2478 Ω/cm^2^) of Calu-3 cell cultures reached statistical significance compared to earlier time points (Fig. 1A). Furthermore, the culture reached maximum polarization, as evidenced by the similar TEER values obtained at 12, 16, and 20 days post-seeding (Fig. 1A).

**Figure 1.**
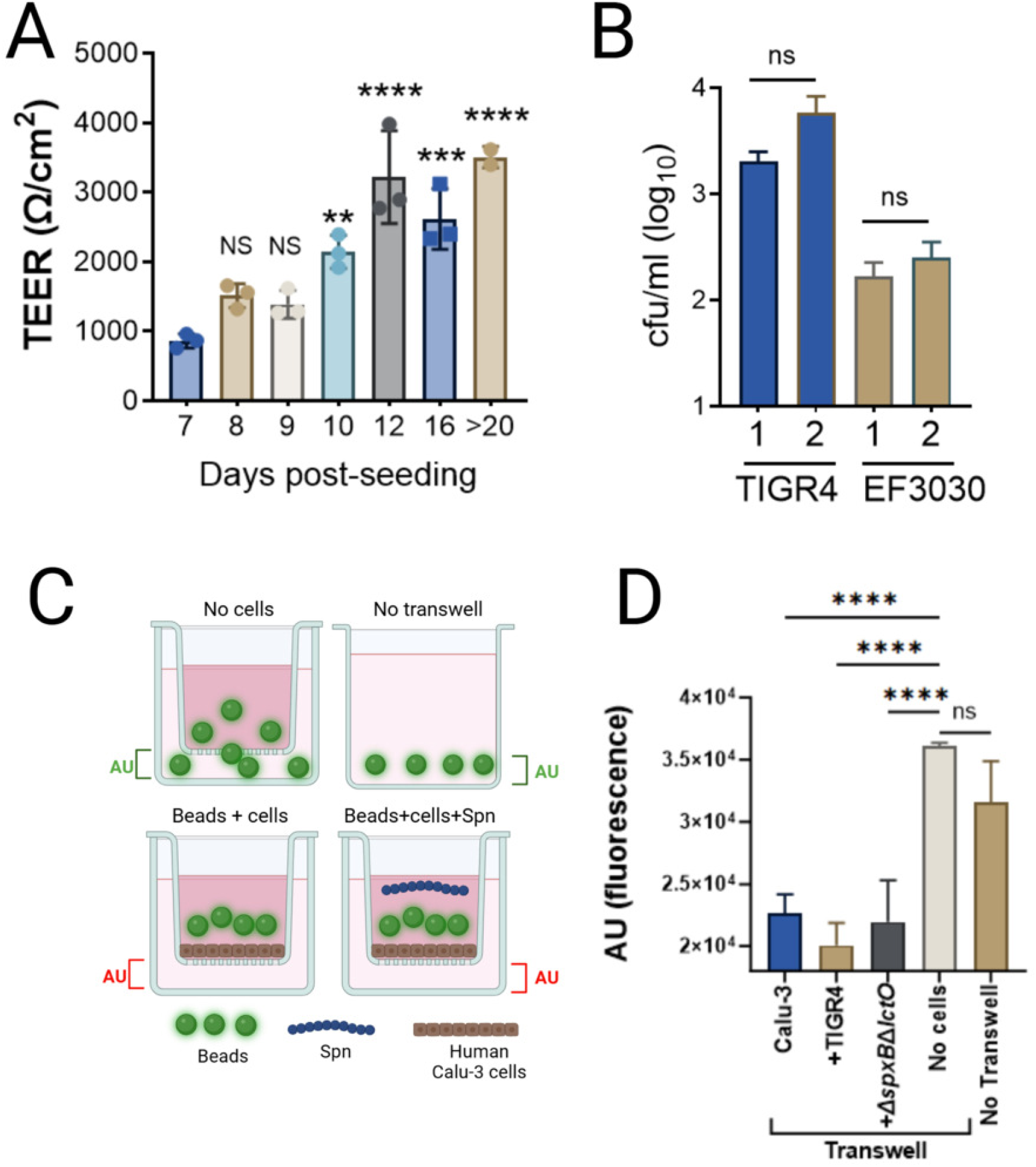
Pneumococcal invasion and translocation across polarized human lung epithelial monolayers. (A) Transepithelial electrical resistance (TEER) of Calu-3 cell monolayers grown on Transwell inserts was measured on the indicated days post-seeding. (B) Polarized Calu-3 lung epithelial cells on Transwell inserts were infected with *S. pneumoniae* strains TIGR4 or EF3030. Translocation of pneumococci to the basolateral compartment was quantified at 1 and 2 h post-infection by plating serial dilutions of basolateral samples and enumerating colony-forming units (CFU/ml). (C) Schematic representation of the Transwell system with or without polarized cells, illustrating the experimental conditions used in panel D. (D) Polarized Calu-3 monolayers on Transwell inserts, empty wells without Transwell devices, or Transwell devices without cells were infected (or left uninfected) with *S. pneumoniae* strains and incubated with fluorescent beads for 2 h. Fluorescence in the basolateral compartment was quantified using a microplate reader (arbitrary units, AU). Data in all panels are presented as mean ± standard error (SE) from at least three independent biological replicates. Statistical comparisons of TEER (A), bacterial density (B), or fluorescence intensity (D) were performed using Student’s t test or one-way ANOVA with Dunnett’s multiple-comparisons test. ns, not significant; **p < 0.005; ***p < 0.0005; ****p < 0.0001.

Polarized bronchial Calu-3 cells were then infected with *S. pneumoniae* strain TIGR4^14^, or EF3030^15^, and infected cells were incubated for 1 or 2 h. Culture medium from the bottom compartment of the Transwell device was collected and pneumococci that had translocated to the basolateral side were counted. Pneumococci translocated after only 1 h post-infection (Fig. 1B). At 2 h post-infection, the density of translocated wild-type bacteria remained similar compared to the 1 h incubation period (*p*>0.156)(Fig. 1B).

Due to the rapid translocation of pneumococci, we conducted further assessments of the integrity of the bronchial cell monolayer. This involved incubating fluorescent beads (diameter 0.2 μm) in our transwell system and collecting fluorescence intensity from the bottom compartment (Fig. 1C). We reasoned that if the monolayer was not confluent, or if pneumococci had caused the detachment of bronchial cells, the beads would be able to traverse the transwell membrane (1.0 μm), resulting in an increase in fluorescence intensity in the bottom compartment. In control transwells without bronchial cells, and in wells without a transwell but with added beads, the fluorescence reached >3.05×10^4^ arbitrary units (AU) (Fig. 1D). In contrast, the fluorescence in transwells with polarized bronchial Calu-3 cells was significantly reduced (Fig. 1D). Pneumococcal infection could have also compromised the integrity of the bronchial cell monolayer. To investigate this, polarized cultures of Calu-3 cells were infected with TIGR4 or TIGR4Δ*spxB*Δ*lctO* (as explained later), then exposed to fluorescent beads and incubated for 2 h. The fluorescence intensity in infected Calu-3 cell cultures remained as low as in the uninfected control (Fig. 1D).

### Disruption of the Pyruvate Metabolic Node Enhances Pneumococcal Translocation Across Polarized Human Lung Epithelial Monolayers

Previous studies have linked the autolysin LytA to increased pneumococcal lung invasion through its role in capsule shedding^11^. Our recent findings revealed that pneumococci lacking the pyruvate node enzymes pyruvate oxidase (SpxB) and lactate oxidase (LctO) invaded bronchial and alveolar cells more rapidly than the wild-type strain^16^. To investigate the role of the pyruvate node enzymes in pneumococcal invasion, we assessed invasion of polarized Calu-3 cells by strain TIGR4Δ*spxB*Δ*lctO* and as a control we used TIGR4Δ*lytA*, lacking pyruvate node enzymes and the major autolysin LytA, respectively. Since TIGR4Δ*spxB*Δ*lctO* lacks production of capsule^17,18^, we also assessed a capsule-knockout TIGR4Δ*cps* strain, which produces wild-type levels of pyruvate oxidase and lactate oxidase^18^.

The translocation of TIGR4Δ*lytA* to the basolateral side was ∼10-fold reduced at both 1 h and 2 h post-inoculation (median, 3.6×10^2^ cfu/ml) but it did not reach statistical significance (Fig. 2A, and 2B). Notably, infection with TIGR4Δ*spxB*Δ*lctO* resulted in a significant increase in translocated pneumococci 2 h post-infection (Fig. 2B). Translocation through Calu-3 cells by TIGR4Δ*cps* was comparable to that of TIGR4 (data not shown). These findings suggest that the capsule, while a crucial virulence factor, does not significantly contribute to the enhanced translocation observed in the mutant deficient in the pyruvate node enzymes. Similarly, translocation of TIGR4Δ*spxB*Δ*lctO* through human alveolar A549 cells and pharyngeal Detroit 562 cells was significantly higher compared to that of the wild-type strain TIGR4 (Fig. 2D and not shown). We further assessed an additional Spn strain, EF3030, to determine whether the observed phenotype was conserved across strains. The EF3030Δ*spxB*Δ*lctO* mutant exhibited significantly higher transcellular than the parent wild-type strain (Fig. 3A). Consistent with these findings, strains lacking pyruvate node enzymes also displayed increased translocation across polarized epithelial cells in an air-liquid interface model (Fig. 3B and 3C).

**Figure 2.**
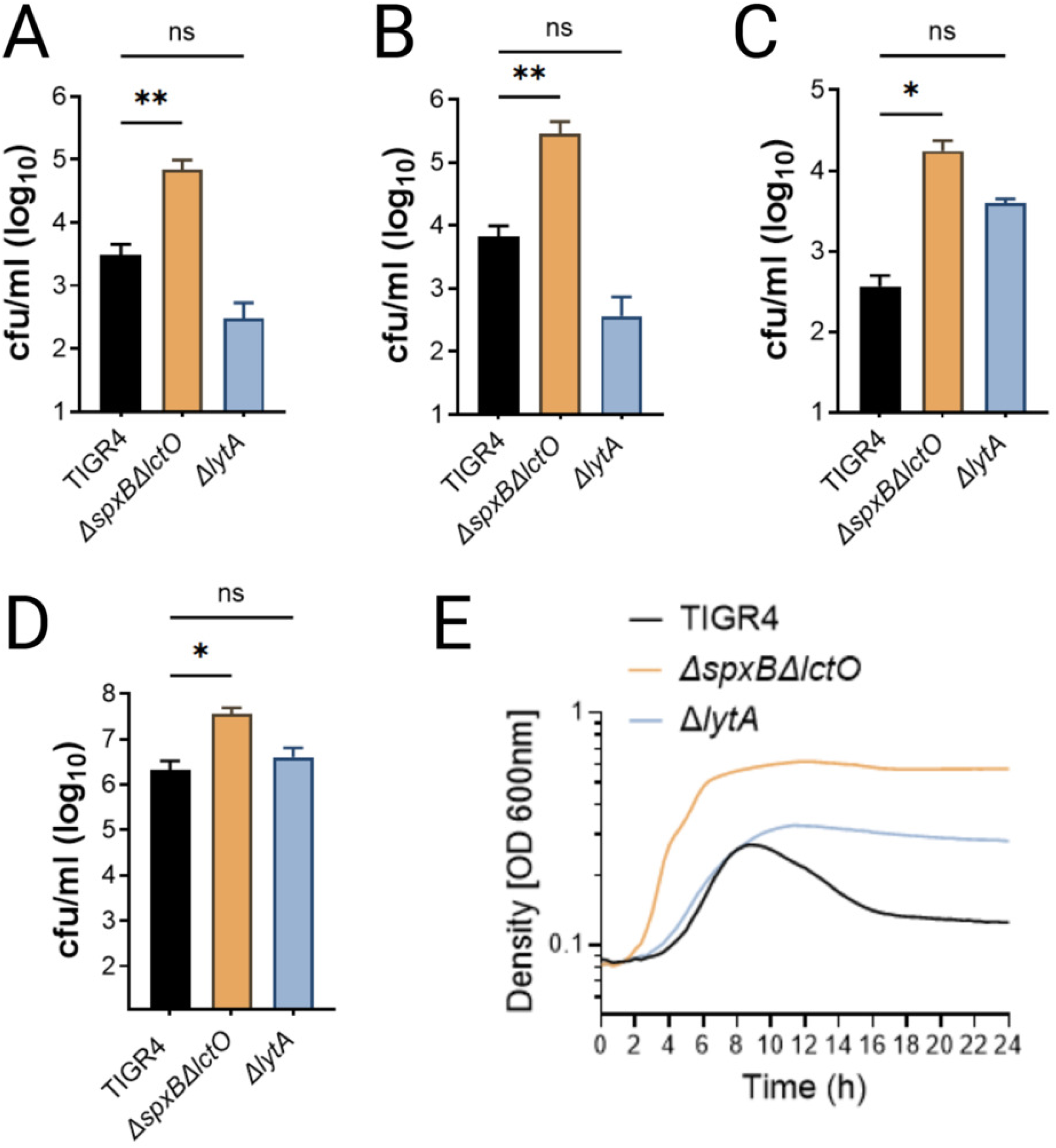
Enhanced adherence and translocation of pyruvate node-deficient pneumococci across human respiratory epithelial monolayers. (A–D) Polarized human lung epithelial cells grown on Transwell inserts were infected with *S. pneumoniae* strains TIGR4, TIGR4 Δ*spxB*Δ*lctO*, or TIGR4Δ*lytA*. Translocation of pneumococci to the basolateral compartment was quantified at 1 h (A) and 2 h (B) post-infection in Calu-3 cells and at 2 h post-infection in A549 cells (C). (D) Pneumococci adherent to Calu-3 cells were quantified after 2 h of incubation. (E) Growth curves of the indicated strains cultured in THY broth at 37°C under 21% O₂ and 5% CO₂. Optical density at 600 nm (OD₆₀₀) was measured every 20 min for 24 h. Data in all panels are presented as mean ± standard error (SE). Statistical comparisons were performed using one-way ANOVA with Dunnett’s multiple-comparisons test (each mutant strain compared to the TIGR4 wild-type control). ns, not significant; *p < 0.05; **p < 0.005.

**Figure 3.**
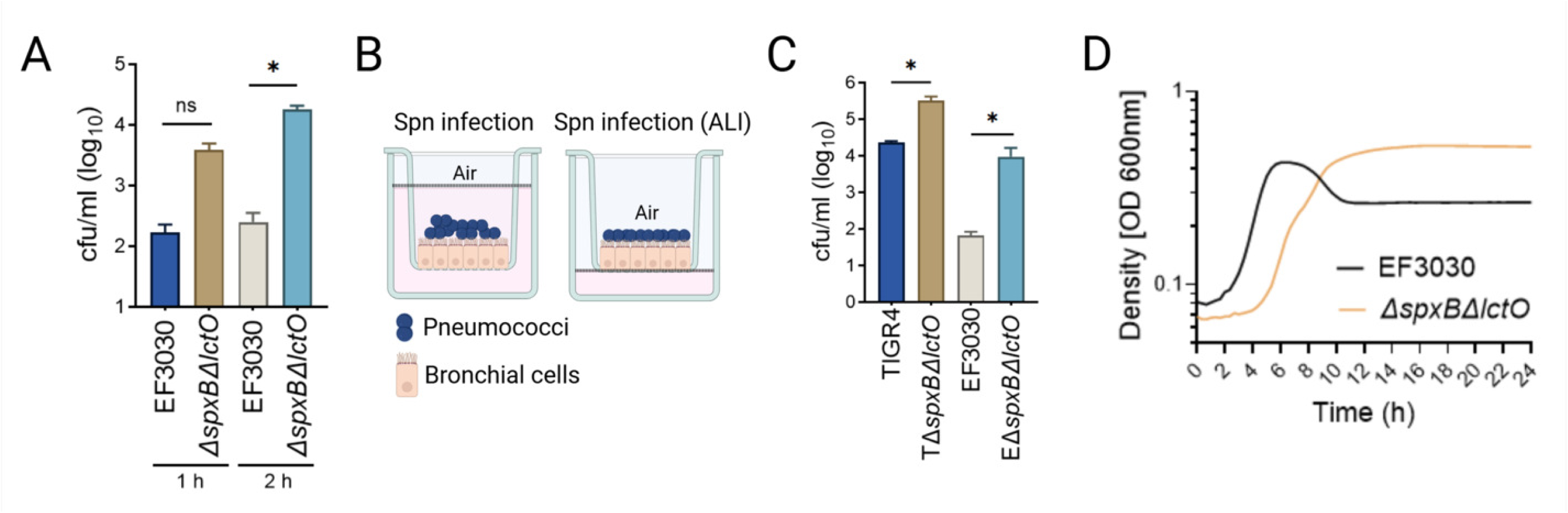
Reproducible pyruvate node-dependent invasion phenotype across pneumococcal strains in human respiratory epithelial monolayers. (A) Polarized human lung Calu-3 cells grown on Transwell inserts were infected apically with *S. pneumoniae* strain EF3030 or its isogenic double mutant EF3030Δ*spxB*Δ*lctO*. Translocation of pneumococci to the basolateral compartment was quantified at 1 h and 2 h post-infection (cfu/ml). (B) Schematic diagram of polarized bronchial epithelial cells cultured in Transwell inserts under standard submerged conditions (left) or air-liquid interface (ALI) conditions (right), illustrating apical infection with pneumococci. (C) Polarized Calu-3 cells grown under ALI conditions were infected apically with the indicated strains (TIGR4, TIGR4Δ*spxB*Δ*lctO*, EF3030, or EF3030Δ*spxB*Δ*lctO*) for 2 h. Translocation of pneumococci to the basolateral compartment was quantified (cfu/ml). (D) Growth curves of wild-type EF3030 and its isogenic double mutant Δ*spxB*Δ*lctO* cultured in THY broth at 37°C under 21% O₂ and 5% CO₂. Optical density at 600 nm (OD₆₀₀) was measured every 20 min for 24 h. Data in panels (A), (C), and (D) represent the mean of at least three independent experiments. Data in panels (A) and (C) are shown as mean ± standard error (SE). Statistical comparisons were performed using Student’s t-test (mutant strains versus the respective parental wild-type strain under each condition). ns, not significant; p < 0.05.

Growth curve analyses confirmed that none of the mutant strains examined showed significant differences in planktonic growth relative to their corresponding wild-type parental strains during the 2h infection period (Fig. 2E and 3D). In parallel, this growth curve analysis verified the expected autolysis defect resulting from the absence of SpxB activity^19^.

We further investigated the adherence of these TIGR4-derivative strains to Calu-3 cells. After a 2 h infection period, TIGR4Δ*spxB*Δ*lctO* exhibited significantly greater adherence to human bronchial cells compared to TIGR4 or TIGR4Δ*lytA* (Fig. 2C). Subsequently, we calculated the translocation frequency, defined as the ratio of adherent bacteria to translocated pneumococci. Among the adherent bacteria, the proportion of pneumococci that translocated to the basolateral side was 1.5×10^−1^ for TIGR4, 8.0×10^−1^ for TIGR4Δ*spxB*Δ*lctO*, and 9.1×10^−3^ for TIGR4Δ*lytA*. Notably, the ratio of adherent to translocated bacteria indicated an >8-fold increase in translocation in pneumococci with a deficiency in pyruvate node enzymes SpxB and LctO compared with those producing these enzymes. Collectively, our findings demonstrate that mutations in *spxB* and *lctO* result in enhanced adherence and translocation of pneumococci.

### The enhanced invasion observed in pneumococci deficient in pyruvate node enzymes is not attributable to host cell toxicity but rather reflects an active translocation mechanism

To investigate whether the enhanced translocation phenotype is caused by cytotoxicity, we monitored transepithelial electrical resistance (TEER) as a sensitive indicator of tight junction integrity. TEER values remained stable throughout the 2 h period of pneumococcal translocation to the basolateral compartment (Fig. 4A). In contrast, a significant decline in TEER was observed in Calu-3 monolayers infected with Spn strains and incubated for 16 h (Fig. 4A).

**Figure 4.**
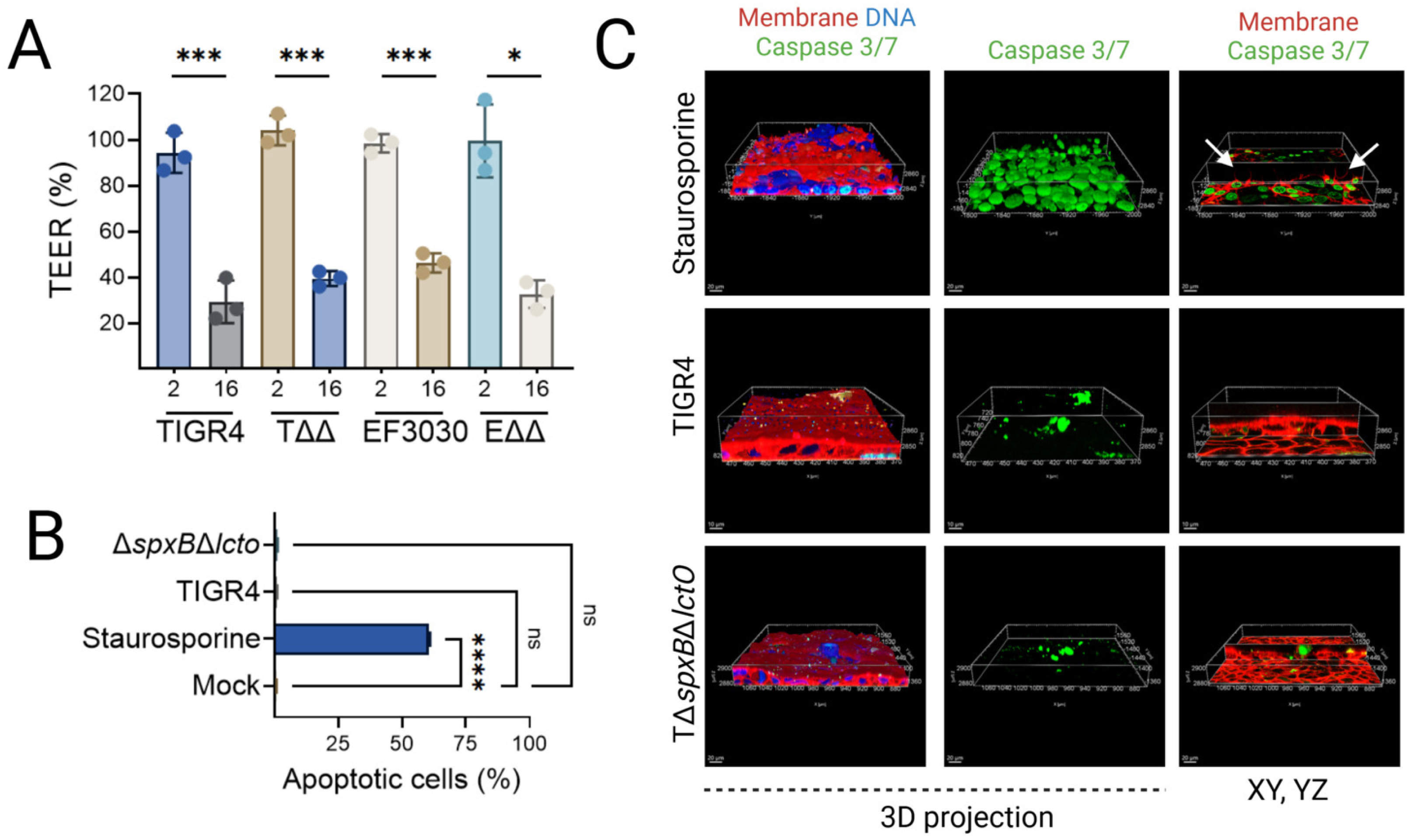
Limited contribution of cytotoxicity to the pyruvate node-dependent invasion phenotype in human respiratory epithelial monolayers. (A) Polarized human lung Calu-3 cells grown on Transwell inserts were infected apically with *S. pneumoniae* strains TIGR4, TIGR4Δ*spxB*Δ*lctO* (TΔΔ), EF3030, or EF3030Δ*spxB*Δ*lctO* (EΔΔ). Transepithelial electrical resistance (TEER) was measured at 2 h and 16 h post-infection and expressed as a percentage of the pre-infection value. (B) Calu-3 cells were mock-infected, infected with TIGR4 or TIGR4Δ*spxB*Δ*lctO* (MOI as in invasion assays), or treated with staurosporine (10 µM) for 4 h. Cells were stained with CellEvent Caspase– 3/7 Green Detection Reagent and SYTOX Blue Dead Cell Stain, followed by flow cytometric analysis. Data show the percentage of apoptotic cells (caspase-3/7-positive, SYTOX Blue-negative). (C) Calu-3 cells were treated with staurosporine (10 µM, 4 h) or infected with TIGR4 or TIGR4Δ*spxB*Δ*lctO* for 4 h, then stained with wheat germ agglutinin (WGA; membrane, red), CellEvent Caspase-3/7 Green Detection Reagent (green), and DAPI (DNA, blue). Z-stack images were acquired by confocal microscopy. Representative 3D projections (left) and corresponding XY and YZ orthogonal sections (right) are shown. Data in panels (A) and (B) represent the mean ± standard error of the mean (SEM) from two independent experiments, each performed with four (A) or two (B) technical replicates. Statistical analysis was performed by one-way ANOVA with Dunnett’s multiple-comparison post hoc test. ns, not significant; p < 0.05; *p < 0.01; **p < 0.001; ***p < 0.0001.

Spn infection is known to activate both apoptotic and pyroptotic pathways in various cell culture models, underscoring complex cytotoxic mechanisms^20–23^. We therefore assessed whether the TIGR4Δ*spxB*Δ*lctO* double mutant triggered apoptosis or pyroptosis that could contribute to its enhanced invasive phenotype. Caspase-3/7 activation, a hallmark of apoptosis, was quantified by flow cytometry and visualized by live high-resolution confocal microscopy. No caspase-3/7 activity was detected in mock-infected cells (Fig. 4B). As expected, staurosporine, a potent apoptosis inducer, elicited robust caspase-3/7 activation after 4 h of treatment; this time point was selected for comparison because it was required to observe clear caspase-3/7 activation with the positive control. In contrast, neither the wild-type TIGR4 strain nor the TIGR4Δ*spxB*Δ*lctO* mutant induced detectable caspase-3/7 activation in lung epithelial cells at this time point (Fig. 4B). Live confocal microscopy corroborated these findings, revealing characteristic caspase-3/7-positive nuclei and membrane blebbing exclusively in staurosporine-treated cells (Fig. 4C, arrows), whereas minimal signal was observed in cells infected with either Spn strain (Fig. 4C).

Infection of lung epithelial cells with Spn for periods exceeding >4 h leads to the secretion of mature IL-18, consistent with the activation of the NLRP3 inflammasome and associated cell death^21,24^. Supernatants from uninfected human bronchial Calu-3 cells or cells infected with TIGR4 for 2 h contained negligible levels of mature IL-18 (<6 pg/mg of total protein). In contrast, supernatants from Calu-3 cells infected with TIGR4 for 6 h contained markedly higher concentrations of mature IL-18 (110 pg/mg) (not shown). Collectively, these data demonstrate that the rapid transcellular migration of pyruvate node mutants occurs independently of host cell cytotoxicity or barrier disruption during the early phase of infection.

### Extracellular Hydrogen Peroxide Does Not Restrict Pneumococcal Transcellular Migration

Previous studies have demonstrated that intracellular pneumococci induce elevated reactive oxygen species (ROS) levels in lung epithelial cells, resulting in disruption of the actin and microtubule cytoskeleton^16,18^. The pyruvate node enzyme SpxB and LctO constitute the primary sources of hydrogen peroxide (H₂O₂) production in (Fig. 5A). Because strains lacking these enzymes (Δ*spxB*Δ*lctO*) exhibit enhanced translocation relative to their wild-type parental strains, we examined whether extracellular H₂O₂ released by bacteria impairs pneumococcal translocation. To test this, Calu-3 monolayers were infected with TIGR4 or EF3030 in medium supplemented with or without catalase (Fig. 5B). Catalase-mediated scavenging of extracellular H₂O₂ did not significantly alter transcellular migration (Fig. 5C) or bacterial adherence (Fig. 5D) for either strain.

**Figure 5.**
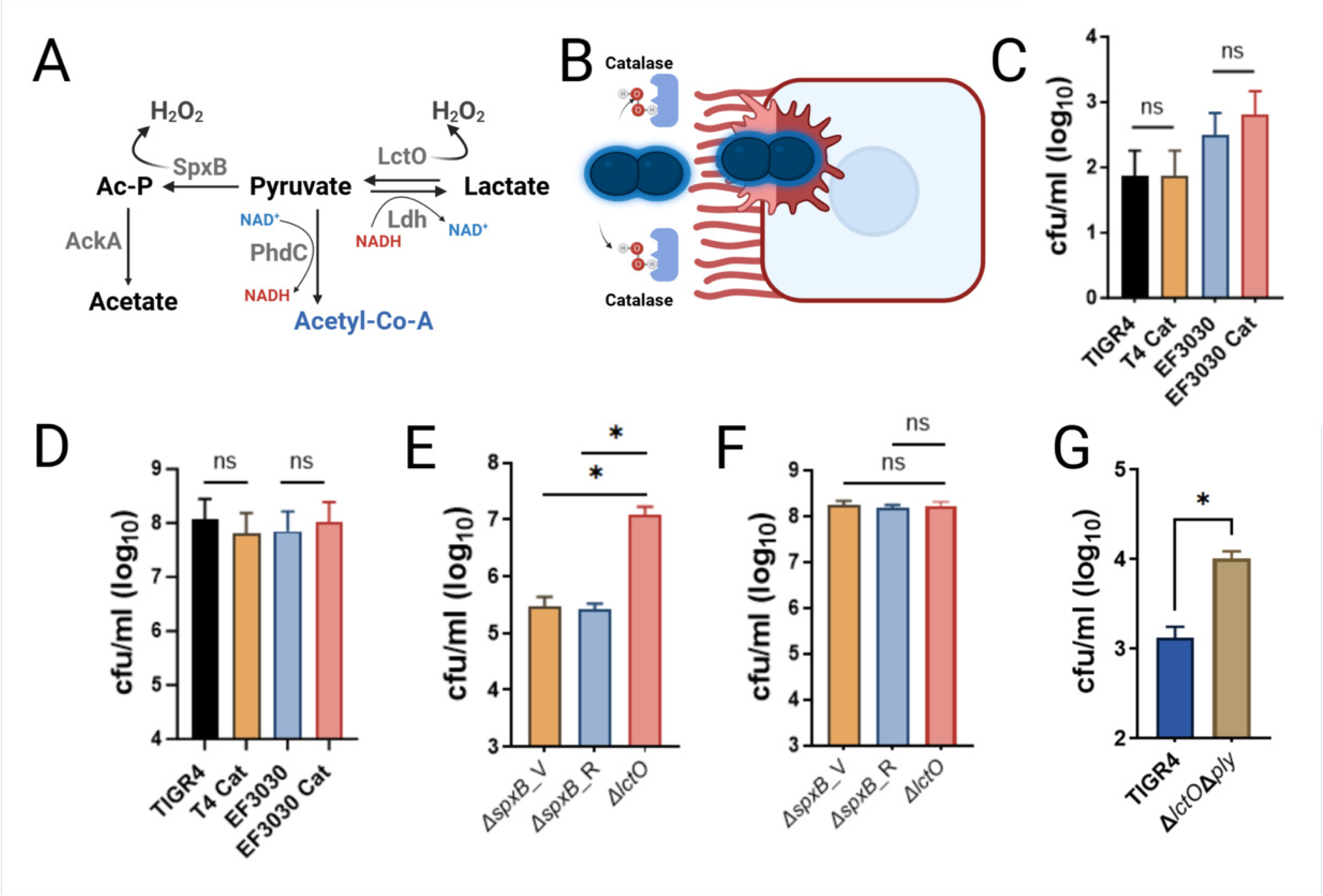
Lactate oxidase (LctO) contributes to the invasion phenotype of *Streptococcus pneumoniae* in lung epithelial cells. (A) Schematic of the pyruvate metabolic node highlighting the key reactions catalyzed by SpxB (pyruvate oxidase) and LctO (lactate oxidase) and their links to central metabolism. (B) Schematic of the Calu-3 Transwell transmigration assay in the presence of extracellular catalase (200 U/mL) to scavenge H₂O₂ produced by pneumococci. (C, D) Effect of catalase supplementation on pneumococcal translocation (C) and adherence (D) across polarized Calu-3 monolayers at 2 h post-infection with strains TIGR4 and EF3030. (E, F) Translocation (E) and adherence (F) of TIGR4 wild-type and its isogenic mutants [TIGR4Δ*spxB* (Δ*spxB*_V), TIGR4 *spxB*^-^ (Δ*spxB*_R), and TIGR4Δ*lctO* (Δ*lctO*)] across polarized Calu-3 cells at 2 h post-infection. (G) Translocation of TIGR4 and the double mutant TIGR4Δ*lctO*Δ*ply* across Calu-3 monolayers at 2 h post-infection. Data represent the mean ± standard error of the mean (SEM) from at least three independent experiments performed in duplicate. Statistical significance was determined using (E and F) one-way ANOVA with Dunnett’s multiple-comparisons test (each Δ*spxB* mutant strain compared to the Δ*lctO* mutant) or (C, D, and G) Student’s t-test. ns, not significant; \**p* ≤ 0.01.

### The absence of lactate oxidase activity enhances the translocation of pneumococci through human respiratory cells

Having excluded passive mechanisms such as tight junction disruption or extracellular H₂O₂ toxicity, we next investigated whether bacterial-host metabolic crosstalk during short-term infection could explain the invasive phenotype. To directly compare the contributions of the two major oxidases, we evaluated the transmigration capacities of isogenic TIGR4Δ*spxB* (lacking pyruvate oxidase) and TIGR4Δ*lctO* (lacking lactate oxidase) mutants.

Notably, TIGR4Δ*lctO* mutants exhibited significantly greater translocation through Calu-3 cells at 2 h post-infection compared to TIGR4Δ*spxB* mutants (Fig. 5E), despite comparable adherence levels (Fig. 5F). This enhanced transmigration phenotype was reproducibly observed using an independent *lctO* mutant strain (TIGR4 *lctO*⁻) and was also evident in additional respiratory cell lines, including alveolar A549 and pharyngeal Detroit 562 cells (not shown). Because pneumolysin (Ply) is known to associate with the pneumococcal cell wall and promote bacterial aggregation^25,26^, we tested whether Ply-mediated aggregation contributes to the enhanced transmigration observed in the Δ*lctO* background. Construction of a *ply* deletion in the TIGR4Δ*lctO* strain showed that the double mutant (TIGR4Δ*lctO*Δ*ply*) displayed transmigration and adherence phenotypes indistinguishable from those of the single Δ*lctO* mutant (Fig. 5G).

### Macrophages phagocytose pneumococci deficient in lactate oxidase at a rate comparable to wild-type pneumococci

To determine if the enhanced epithelial cell invasion observed in lactate oxidase-deficient pneumococci extends to macrophages and dendritic cells, we infected murine IC21 macrophages and JAWS II dendritic cells with TIGR4, TIGR4Δ*spxB,* and TIGR4Δ*lctO*. After 30 min, phagocytosis of TIGR4 and TIGR4Δ*lctO* was comparable, while TIGR4Δ*spxB,* with reduced capsule expression^10,17^, exhibited significantly increased phagocytosis (Fig. 6A and 6B). These findings suggest that capsule polysaccharide-mediated resistance to phagocytosis remains intact in lactate oxidase-deficient pneumococci. Notably, murine macrophages and dendritic cells infected with TIGR4Δ*lctO* displayed increased toxicity compared to the wild-type TIGR4 strain (Fig. 6C and 6D). To test whether this enhanced toxicity was mediated by increased pneumolysin (Ply) activity, we tested a TIGR4 Δ*lctO*Δ*ply* double mutant. Phagocytosis of the double mutant was similar to that of TIGR4Δ*lctO* (Fig. 6A and 6B); however, deletion of *ply* completely abrogated the toxicity observed with the single Δ*lctO* mutant (Fig. 6C and 6D). These findings demonstrate that, under the conditions of this macrophage and dendritic cell phagocytosis assay, pneumolysin plays a critical role in the enhanced toxicity elicited by lactate oxidase-deficient pneumococci.

**Figure 6.**
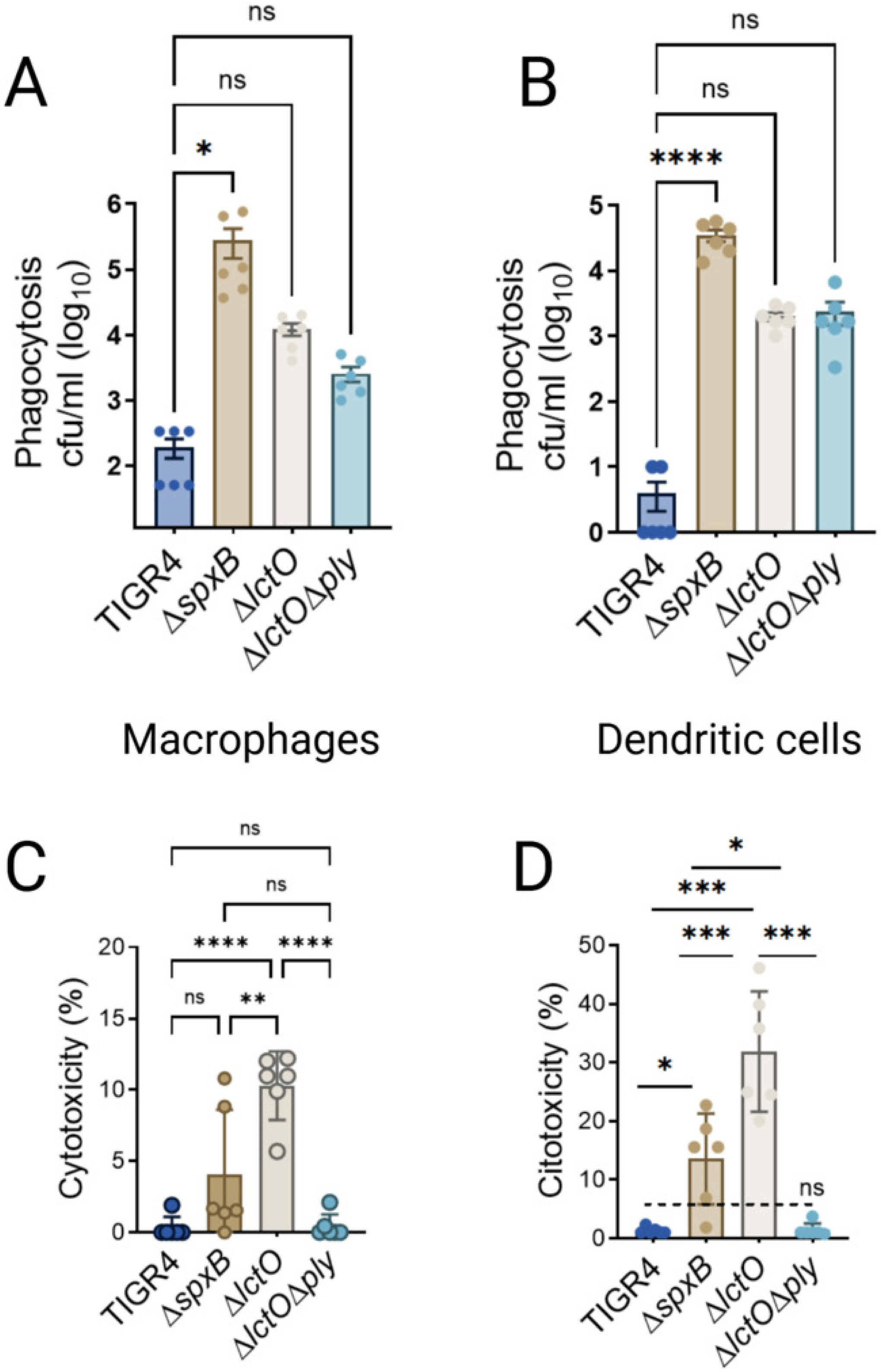
Lactate oxidase (LctO) does not affect phagocytosis of *Streptococcus pneumoniae* by macrophages and dendritic cells. (A, B) Phagocytosis of TIGR4, TIGR4Δ*spxB*, TIGR4Δ*lctO*, and TIGR4Δ*lctO*Δ*ply* by IC21 macrophages (A), or JAWSII-dendritic cells (B), 30 min post-infection. (C, D) Cytotoxicity of the indicated strains toward IC21 macrophages (C), or JAWSII-dendritic cells (D), measured at 24 h post-infection. Data represent the mean ± standard error of the mean (SEM) from three independent experiments performed in duplicate. Statistical significance was determined using one-way ANOVA with Dunnett’s multiple-comparisons test. ns, not significant; *p ≤ 0.05; **p=0.0029; **p=0.0005; ****p ≤ 0.0001.

### Intracellular pneumococci are located into membrane-bound vesicles

To gain further insights into the molecular mechanisms underlying pneumococcal translocation, human alveolar or bronchial Calu-3 cells were infected with wild-type strains or their corresponding mutant derivatives. Subsequently, a series of cellular and bacterial components were stained, including cellular actin, N-acetyl-D-glucosamine (GlcNAc)-, sialic acid (SA)-containing membranes, and DNA. These samples were then analyzed using high-resolution confocal microscopy. Microscopic examination of the optical section from cells adhered to the substrate (i.e., bottom) revealed a typical actin cytoskeleton characterized by stress fibers in both mock-infected, TIGR4Δ*lctO*-infected, or EF3030Δ*lctO*-infected (i.e., SpnΔ*lctO*) human bronchial cells after 1 h of incubation (Fig. 7A). No pneumococci were observed intracellularly in cells infected with wild-type strains TIGR4 or EF3030 at this time point (not shown). A mid-plane optical section of SpnΔ*lctO*-infected cells revealed intracellular bacteria in close proximity to nuclei as soon as 1 h post-infection (Fig. 7A, arrows). Strikingly, while the GlcNAc/SA signal was virtually undetectable in mid-plane optical sections of mock-infected cells, as expected given its integral role in the eukaryotic cell membrane, a pronounced intracellular GlcNAc/SA signal was observed in SpnΔ*lctO*-infected cells, specifically localized around the invading bacteria (Fig. 7A, arrowheads). XZ and YZ optical sections confirmed the presence of the GlcNAc/SA signal in the membrane of mock-infected cells, but both membrane and intracellular signals of GlcNAc/SA were observed in SpnΔ*lctO*-infected bronchial cells (Fig. 7B).

**Figure 7.**
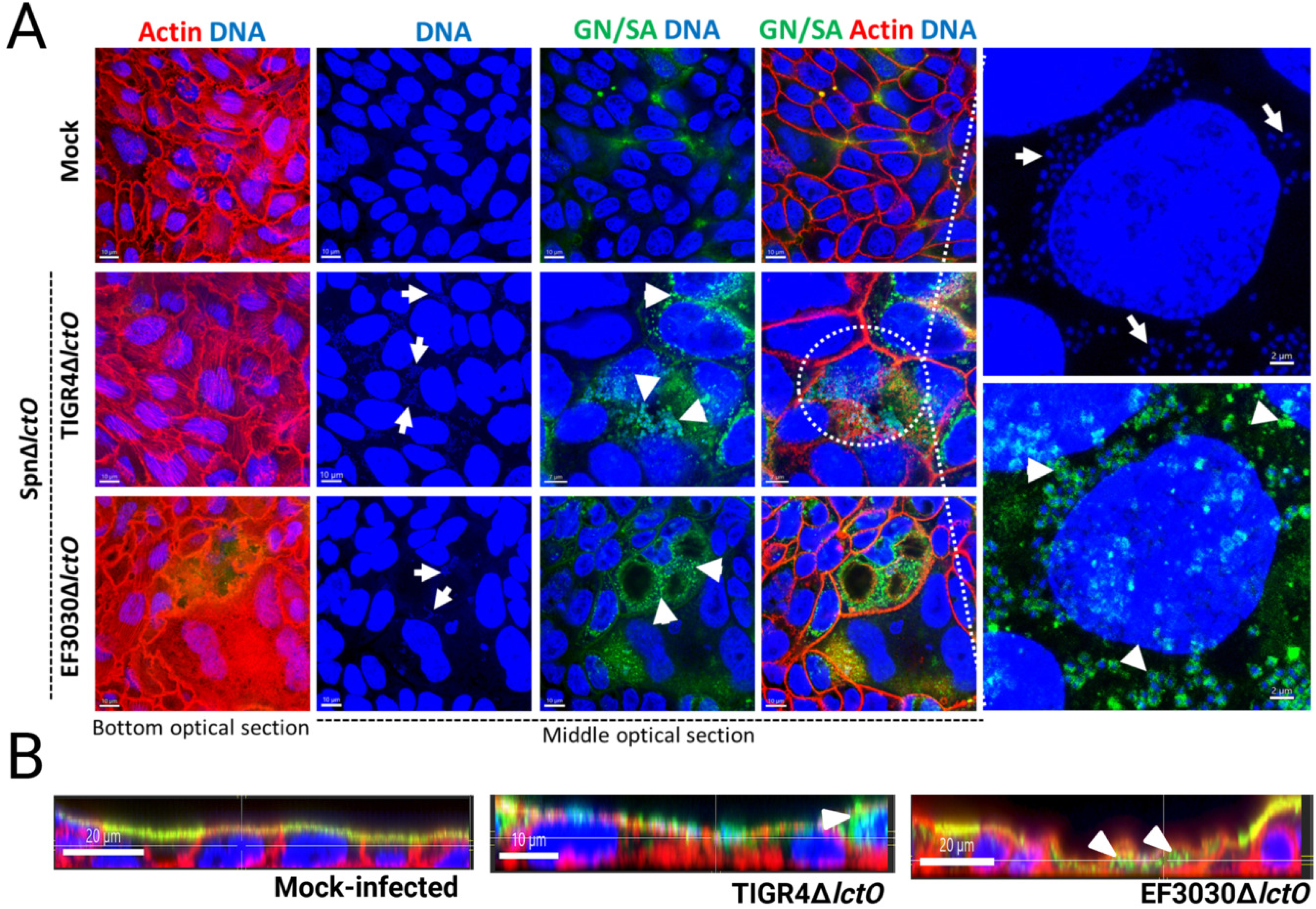
High-resolution confocal imaging of enhanced invasion of lung epithelial cells by pneumococci lacking lactate oxidase (LctO). (A) Polarized human lung Calu-3 cells grown on eight-well chamber slides were mock-infected or infected with *S. pneumoniae* strains TIGR4Δ*lctO* or EF3030Δ*lctO* for 1 h. After fixation with 2% paraformaldehyde and permeabilization, cells were stained for actin (red), DNA (blue), and N-acetylglucosamine/sialic acid (GN/SA, green) using wheat germ agglutinin (WGA). Representative confocal images of the bottom optical section and the middle optical section are shown. Arrows indicate intracellular pneumococci colocalizing with intracellular GN/SA signal; arrowheads mark additional intracellular bacteria. (B) Orthogonal XZ views from 3D reconstructions confirming the intracellular localization of pneumococci within GN/SA-containing compartments (arrowheads) in Δ*lctO* mutant-infected monolayers.

Collectively, these data demonstrate that ablation of LctO activity—independent of H₂O₂ production (via SpxB), pneumolysin function, or strain background—consistently augments pneumococcal transmigration across diverse human respiratory epithelial barriers. These findings implicate LctO-mediated lactate-to-pyruvate oxidation as a critical negative regulator of pneumococcal invasion at the epithelial interface and, by extension, of subsequent lung pathogenesis.

## Discussion

In this study, we demonstrate that *Streptococcus pneumoniae* rapidly translocates across polarized lung epithelial barriers and that this process is regulated by the pyruvate node enzyme lactate oxidase (LctO). Deletion of *lctO*, alone or in combination with *spxB*, significantly enhances transmigration across human bronchial (Calu-3), alveolar (A549), and pharyngeal (Detroit 562) epithelial monolayers. The hyper-invasive phenotype was reproducible across strains (TIGR4 and EF3030) and culture conditions, including air-liquid interface models. Notably, the Δ*lctO* single mutant consistently showed stronger transmigration than the Δ*spxB* mutant, indicating a dominant role for LctO in restraining invasion. Consistent with our *in vitro* findings, a previous study reported an increased bacterial burden in the blood of mice following intranasal infection with a TIGR4Δ*lctO* mutant compared with the isogenic wild-type strain^27^.

The enhanced translocation of pyruvate node mutants occurred without early disruption of epithelial barrier integrity, as evidenced by stable transepithelial electrical resistance (TEER) during the first 2 h of infection. Consistent with this, Δ*spxB*Δ*lctO* mutants did not induce detectable caspase-3/7 activation or substantial IL-18 release at early time points, in contrast to later stages of infection or staurosporine treatment. These results align with our recent observations that infection with wild-type TIGR4 or EF3030 strains does not trigger caspase-3/7 activation in human A549 lung epithelial cells even after up to 10 h under these culture conditions^28^. However, animal models and *ex vivo* studies of pneumococcal pneumonia have documented multiple cell death markers consistent with the simultaneous activation of apoptosis and pyroptosis^20–23^ indicating that the mechanisms underlying lung cytotoxicity are complex and time-dependent. High-resolution confocal microscopy further revealed that Δ*lctO* mutants were internalized and localized within N-acetylglucosamine/sialic acid (GN/SA)-containing intracellular compartments as early as 1 h post-infection, whereas wild-type bacteria were rarely observed intracellularly at this time point. Together, these data strongly support an active transcellular migration mechanism rather than passive barrier disruption.

The increased transmigration of Δ*lctO* mutants correlated with enhanced adherence to epithelial cells but was independent of capsule expression, extracellular H₂O₂, and pneumolysin function, as shown by experiments with catalase supplementation, capsule mutants, and the Δ*lctO*Δ*ply* double mutant. In murine IC21 macrophages and JAWS II dendritic cells, Δ*lctO* mutants were phagocytosed at rates comparable to wild-type TIGR4, although they induced greater pneumolysin-dependent cytotoxicity at 24 h. Collectively, these data identify LctO as a critical negative regulator of pneumococcal invasion at the respiratory epithelial interface.

The pyruvate-node, coupled to the glycolytic pathway, has previously been implicated in capsule production in certain *S. pneumoniae* strains^17,19^. Building on prior work showing rapid capsule shedding upon epithelial contact^10,11^, our findings indicate that the enhanced transmigration of Δ*lctO* mutants is largely independent of capsule status, as deletion of *lytA* or the capsule locus did not recapitulate the phenotype. The pronounced intracellular redistribution of GN/SA signal surrounding invading Δ*lctO* bacteria suggests that these mutants may traffic more efficiently through or modify endocytic compartments. These observations are consistent with reports that certain pneumococcal serotype 3 strains recruit GN/SA-containing membrane components *in vivo*, potentially to evade host immune recognition^29^. We hypothesize that LctO activity in wild-type pneumococci limits the recruitment or intracellular redistribution of host GN/SA-containing membranes, thereby restraining bacterial invasion. In support of this, a recent study demonstrated that Spn might prefers GN instead of glucose when grown under infection-mimicking conditions^30^. This process may represent a species-specific metabolic strategy that links central carbon metabolism with modulation of host membrane trafficking to balance colonization and invasion. Such a mechanism would be consistent with the known role of sialic acid residues as self-associated molecular patterns recognized by mammalian siglecs and other immune receptors^31^. Future studies will be necessary to test this hypothesis and to define the precise molecular links between LctO-dependent redox balance and host endocytic pathways.

This work extends the known roles of lactate-metabolizing enzymes in promoting invasive virulence across Gram-positive pathogens, including *Streptococcus pyogenes*^32,33^ and *Staphylococcus aureus*^34,35^, and demonstrates that modulation of the pyruvate node can shift Spn toward a more invasive phenotype without compromising early epithelial barrier integrity. Moreover, our findings are consistent with emerging evidence that pneumococcal infection induces reprogramming of host glycolysis^28,36^ and that the complex metabolic networks of the pyruvate node play central roles in pneumococcal pathogenesis^10,27,36^. By suppressing lactate efflux and increasing intracellular pyruvate availability, this metabolic crosstalk likely generates a supportive microenvironment that sustains bacterial fermentation and helps mitigate redox stress during the early phases of infection. Collectively, these processes highlight population heterogeneity as a key driver of pneumococcal disease progression, facilitating phenotypic switching from controlled airway colonization to enhanced epithelial invasion and systemic dissemination.

In summary, these results position LctO as a critical metabolic checkpoint that restrains pneumococcal invasion of respiratory epithelia. Further studies are warranted to define the precise redox and metabolic changes underlying the hyper-invasive phenotype of *lctO* mutants and to evaluate the therapeutic potential of targeting this pathway to limit invasive pneumococcal disease.

## Materials and Methods

### Bacterial strains and media

*S. pneumoniae* strains utilized in this study are listed in Table 1. Strains were cultured on blood agar plates (BAP) containing 5% sheep blood from frozen stocks prepared in medium with skim milk-tryptone-glucose-glycerol (STGG^37^) medium. Broth cultures were grown in Todd-Hewitt broth supplemented with 0.5% (w/v) yeast extract (THY). All cell infection experiments were performed using infection medium consisting of either DMEM supplemented with 5% fetal bovine serum (FBS) (Corning), 2 mM L-glutamine (Gibco), and 1X HEPES (Gibco) or EMEM infection medium supplemented with 5% fetal bovine serum (FBS) (Corning), and 1X HEPES (Gibco).

**Table 1.** Strains utilized in this study.

| Strain | Description | Reference or Source |
| --- | --- | --- |
| TIGR4 | Invasive clinical isolate, capsular serotype 4, sensitive to antibiotics | 14 |
| TIGR4Δ*spxB* (SPJV29) | TIGR4 derivative with insertionally inactivated *spxB::ermB*, (Ery<sup>r</sup>) erythromycin resistance, hydrogen peroxide deficient strain | 43 |
| TIGR4 *spxB*<sup>-</sup> | TIGR4 derivative with insertionally inactivated *spxB::ermB*, (Ery<sup>r</sup>) erythromycin resistance, (Rosch lab mutant strain) | 17 |
| TIGR4 *lctO*<sup>-</sup> | TIGR4 derivative with insertionally inactivated *lctO::aad9*, (Spc<sup>r</sup>), (Rosch lab mutant strain) | 17 |
| TIGR4Δ*lctO* (SPJV42) | TIGR4 derivative with insertionally inactivated *lctO::aad9*, (Spc<sup>r</sup>), hydrogen peroxide deficient strain | 44 |
| TIGR4Δ*lctO*Δ*ply* (SPJV76) | SPJV42 derivative with insertionally inactivated *ply::cat* (Cm<sup>R</sup>) | This study |
| TIGR4Δ*spxB*Δ*lctO* (SPJV41) | TIGR4 derivative with *spxB* and *lctO* gene deletion by insertion-deletion with erythromycin and spectinomycin cassette, respectively, hydrogen peroxide deficient strain | 44 |
| TIGR4Δ*lytA* (SPJV62) | TIGR4 derivative with *lytA* gene deletion by insertion-deletion with am erythromycin cassette | This study |
| TIGR4Δ*cps* (P2422) | TIGR4 derivative capsule knockout mutant (Kan<sup>r</sup>) | 45 |
| EF3030 | Invasive clinical isolate, capsular serotype 19F, sensitive to antibiotics | 15 |
| EF3030Δ*spxB*Δ*lctO* (SPJV49) | EF3030 derivative with *spxB* and *lctO* gene deletion by insertion-deletion with erythromycin and spectinomycin cassette, respectively, hydrogen peroxide deficient strain | 43 |
| EF3030Δ*lctO* (SPJV51) | EF3030 with an insertionally inactivated *lctO::aad9* (Spc<sup>r</sup>) | 10 |

### Cell cultures of human respiratory cells

Human respiratory epithelial cell lines were used in this study: A549 human type II alveolar epithelial cells (ATCC CCL-185), Calu-3 human bronchial epithelial cells (ATCC HTB-55), and Detroit 562 human pharyngeal epithelial cells (ATCC CCL-198). A549 cells were maintained in Dulbecco’s Modified Eagle’s Medium (DMEM; Gibco) supplemented with 10% (v/v) fetal bovine serum (FBS; Corning), 2 mM L-glutamine (Gibco), and 100 U/mL penicillin-streptomycin (Gibco). Calu-3 cells were cultured in Eagle’s Minimum Essential Medium (EMEM; ATCC) supplemented with 10% (v/v) FBS (Corning) and 100 U/mL penicillin-streptomycin (Gibco). Detroit 562 cells were grown in DMEM (Gibco) supplemented with 10% (v/v) non-heat-inactivated FBS (Corning), 1% (v/v) non-essential amino acids (Sigma), 1% (v/v) L-glutamine (Sigma), 100 U/mL penicillin-streptomycin (Gibco), and 10 mM HEPES (Gibco). All cell lines were incubated at 37°C in a humidified atmosphere containing 5% CO₂. Cells were incubated at 37°C with 5% CO_2_ and supplemented with fresh medium 3 times weekly and passaged to a new flask once weekly or when cells reached ∼100% confluency by trypsinization with Trypsin-EDTA (0.25%), phenol red (Gibco), and seeded in the experiment-specific device.

### Preparation of inoculum

Bacteria from frozen STGG stocks were streaked onto blood agar plates (BAP) (TSA with Sheep blood) and incubated overnight at 37 °C in a 5% CO₂ atmosphere. Bacterial growth was harvested from the plates by washing with sterile phosphate-buffered saline (PBS) at pH 7.4. The resulting suspension was adjusted to a final optical density at 600 nm (OD₆₀₀) of approximately 0.1, corresponding to ∼5.15 × 10⁸ CFU/mL. Aliquots of these suspensions were routinely serially diluted and plated to confirm bacterial concentrations (CFU/mL). Unless otherwise indicated, all *in vitro* experiments described below were inoculated with freshly prepared inocula immediately prior to the start of each experiment.

### Growth curves

THY was inoculated with Spn strains in 24-well plates (Genesee Scientific). Plates were incubated at 37 °C in a 5% CO₂ atmosphere using a microplate reader (Agilent BioTek Synergy H1). Optical density at 600 nm (OD₆₀₀) was measured every 20 min for 24 h to generate bacterial growth curves.

### Infection of human respiratory cells with *S. pneumoniae* strains

Experiments were conducted once cell cultures reached 100% confluence and, where applicable, became polarized (e.g., Calu-3 cells, as indicated by the development of transepithelial electrical resistance; see below), a process that occurred 10–12 days post-seeding. Cells were cultured under the following conditions: 12-well tissue culture–treated plates (Genesee Scientific, GenClone) with Transwell inserts (Corning Costar) containing 0.4-µm pore membranes; 6-well tissue culture–treated plates (Genesee Scientific, GenClone) with cell culture inserts (Genesee Scientific) containing 1-µm pore membranes; 6-well plates with glass coverslips (Chemglass Life Sciences); or 8-well chamber slides (Lab-Tek II Chamber Slide with Cover, NUNC). All cultures were maintained at 37 °C in a 5% CO₂ atmosphere.

Prior to infection, cells were washed three times with PBS and then incubated with the appropriate infection media—DMEM supplemented with 5% FBS, 1% L-glutamine, and 1% HEPES for A549 cells or EMEM supplemented with 5% FBS and 1% HEPES for Calu-3 cells. Cells were subsequently infected with *S. pneumoniae* strains, along with the appropriate controls, using freshly prepared inocula as described above. In some experiments, 0.2 µm Fluoresbrite® YG Carboxylate beads (Polysciences, Inc., Cat. #21636-1) were used.

### Transepithelial electrical resistance (TEER) measurements

Transepithelial electrical resistance (TEER) was measured to assess tight junction integrity and confirm epithelial polarization prior to infection. Calu-3 cells were seeded onto 12-well tissue culture– treated plates (Genesee Scientific, GenClone) fitted with Transwell inserts (Corning Costar) containing 0.4-µm pore membranes. Beginning 7 days post-seeding, TEER was measured using an EVOM volt/ohm meter (World Precision Instruments) equipped with “chopstick” electrodes. Prior to use, electrodes were sterilized with ethanol and subsequently rinsed with the appropriate cell culture medium, which was also used to obtain blank measurements. TEER values for each well were obtained by placing the electrodes on the apical and basolateral sides of the Transwell insert containing a confluent cell monolayer and were calculated according to the manufacturer instructions.

### Air–liquid interface (ALI) model

Calu-3 cells were cultured on Transwell inserts with 1-µm pore size membranes (Genesee Scientific) and maintained under submerged conditions until a confluent monolayer was established. Monolayer formation and barrier integrity were monitored by measuring TEER. To induce differentiation, an air–liquid interface (ALI) was established 20 days post-seeding by removing medium from the apical compartment, while culture medium was supplied exclusively to the basolateral compartment. Cells were maintained at the ALI for an additional 15 days prior to infection experiments.

### Quantification of IL-18

Supernatants from Transwell cultures infected with *S. pneumoniae* for 2, 4, or 6 h were collected. Supernatants were first centrifuged at 100 × g for 5 min to remove detached host cells, then at 21,130 × g for 5 min to remove remaining debris and pneumococci. The resulting cell-free supernatants were filter-sterilized and stored at −80 °C until analysis. IL-18 levels were quantified using a Human IL-18 ELISA kit from Invitrogen according to the manufacturer’s instructions.

### Translocation of *S. pneumoniae* through Calu-3 cells in the presence of catalase

Calu-3 cells grown on Transwells were infected with the wild-type Spn strain TIGR4 in the presence of catalase (Sigma), which was added to the infection medium at a final concentration of 200 U/mL. Samples were collected from the apical chamber to assess bacterial adhesion and from the basolateral chamber to assess bacterial translocation. Bacterial densities were determined by serial dilution and plating.

### Confocal microscopy experiments

Calu-3 cells were seeded in 8-well chamber slides and used once at 10 days post-seeding. Prior to infection, cells were washed three times with PBS and incubated with EMEM infection medium. Cells were then infected with Spn and incubated at 37 °C in a 5% CO₂ atmosphere. Following infection, cells were washed with PBS and fixed with 2% PFA for 15 min at room temperature. Fixed cells were washed once with PBS, permeabilized with 0.5% Triton X-100 in PBS for 5 min at room temperature, washed again with PBS, and blocked with 2% bovine serum albumin (BSA) for 30 min at room temperature. Cells were then stained for 1 h with either serotype-specific polyclonal antibody (Statens Serum Institute, Denmark) labeled with Alexa-488 (anti-S4-A488; Molecular Probes) at 40 μg/mL^38^, or with wheat germ agglutinin conjugated to Alexa-488 (WGA; ThermoFisher) at 5 μg/mL, in combination with phalloidin (7 units) in 2% BSA. After staining, cells were washed with PBS, air-dried, and mounted with ProLong Diamond Antifade mounting medium containing DAPI (Molecular Probes). Slides were imaged using a Nikon Eclipse C2 laser scanning confocal system mounted on a Ti-E motorized inverted microscope. Excitation was performed using a solid-state 405/488/561/640 nm laser unit and a CFI Apo 60× oil immersion objective (NA 1.40). Images were collected using a high-sensitivity C2-DU3 Detection Unit with 435/34, 525/50, and 600/50 filters. Confocal images were analyzed using Imaris x64 software version 10.1.0.

### Live cell imaging and time-lapse microscopy

Calu-3 cells were seeded onto collagen-coated 35-mm glass-bottom dishes (MatTek) and cultured to confluence. Prior to infection, treatment, and imaging, cells were washed three times with PBS and incubated in infection medium supplemented with Hoechst (50 μg/mL), wheat germ agglutinin–Alexa Fluor 555 (WGA-A555; 1 μg/mL), and CellEvent Caspase-3/7 Green Detection Reagent (2 μM; ThermoFisher). For apoptosis induction, cells were treated with staurosporine (10 μM; Sigma) for 4 h. For infection experiments, cells were exposed to Spn strains and incubated for 4 h at 37 °C in a humidified chamber (H301-Nikon-TI-S-ER) with 5% CO₂ maintained by an Okolab controller. Live-cell confocal imaging and time-lapse acquisition were performed using a Nikon Eclipse C2 laser scanning confocal microscope with the specifications described above. Image processing and analysis of both static and time-lapse datasets were conducted using Imaris software (x64, version 10.1.0).

### Flow cytometry analysis using caspase-3/7 reagent

To assess apoptosis in human A549 lung epithelial cells infected with Spn, the CellEvent Caspase-3/7 Green Detection Reagent (ThermoFisher) was employed. A549 cells grown in a 6-well plate were washed and treated with infection medium as described above. Cells were then infected with Spn and incubated for 4 h. After incubation, cells were lifted from the wells using 0.25% Trypsin-EDTA, counted, and ∼1×10^6^ cells were used for staining following the manufacturer’s protocol. Briefly, cells were incubated with the CellEvent Caspase-3/7 Green Detection Reagent (2 µM final concentration) for 30 minutes at 37°C prior to analysis. As a control, staurosporine (Sigma, 10 µM, 4 hours) was used as a positive control for caspase-3/7 activation. Subsequently, cells were stained with 1 μM SYTOX Blue Dead Cell Stain (Invitrogen) for dead cells. Flow cytometry was performed on a NovoCyte 3000 (Agilent). Caspase-3/7 activation was detected in the FITC channel (530/30 nm filter) following excitation at 488 nm. SYTOX Blue Dead Cell Stain was excited by the violet 405 nm laser and detected in the 530/30 nm channel. Fluorescence compensation, was performed using FMO and single-color controls. Data from 50,000 events per sample were acquired and analyzed using NovoExpress software (Agilent).

### Construction of a *lytA* deletion mutant in TIGR4

Upstream and downstream flanking regions of the *lytA* gene (SP_1937) were PCR amplified from genomic DNA extracted from TIGR4 using the following primer sets: upstream, JV2023-1 (5′-CCTGTGTGAAATTCTTATCCGCTGGATTCCCAGTTGAGTGTGC-3′) and JV2023-2 (5′-TGACCTTTGCCCTTCTTCCT-3′); downstream, JV2023-3 (5′-ACCTTCCTAATGGCATGTCTGA-3′) and JV2023-4 (5′-AAGGAGCTAAAGAGGTCGGCAGGCTGGGTCAAGTACAAGG-3′). An erythromycin resistance cassette (*ermB*) was amplified from TIGR4Ω*ermB*^39,40^ using primers JV2023-5 (5′-GCCGACCTCTTTAGCTCCTT-3′) and JV2023-6 (5′-AGCGGATAAGAATTTCACACAGG-3′). These three PCR fragments were assembled by PCR splicing, and the final product was purified using a QIAquick PCR Purification Kit (Qiagen) for transformation into competent TIGR4 cells. Putative *lytA* deletion mutants were selected on BAP containing erythromycin (1 µg/mL), and the deletion was screened by PCR using primers JV2023-2 and JV2023-3. To confirm the deletion, a selected TIGR4Δ*lytA* clone was subjected to whole-genome sequencing at the SeqCenter (formerly MiGS Microbial Genome Sequencing Center) using the NextSeq 2000 platform. Paired-end reads were assembled and annotated using the RAPT NCBI tools, with reference to the TIGR4 genome (NC_003028.3). The resulting sequences were deposited in NCBI GenBank (it will provided during revisions).

### Cell cultures of IC21 mouse macrophages and culture conditions

IC21 (TIB-186) is a mouse macrophage cell line, and JAWSII (CRL-3612) is a mouse dendritic cell line; both cell lines was obtained from ATCC^®^. IC21-macrophages were maintained in RPMI (Invitrogen) media supplemented with 10% of FBS (HiClone) and 1% of antibiotic/antimycotic (Invitrogen). JAWSII-dendritic cells were maintained in MEM media supplemented with ribonucleotides (Invitrogen), 10% of FBS, and 1% of antibiotic/antimycotic (Invitrogen). Dendritic cells were matured with 100 ng of LPS from *Escherichia coli* (Sigma-Aldrich). Macrophages (3 x 10^5^) or matured dendritic cells were seeded on 24-well plates (Corning), washed once with PBS (Sigma-Aldrich), and then incubated in appropriate media without antibiotics. Cells were infected with *S. pneumoniae* strains. To synchronize the infection, plates were centrifuged at 1400 RPM for 1 min and then incubated for 30 min. After infection, cells were washed three times with PBS to remove extracellular bacteria; infected cells were returned to the incubator for 30 min (to evaluate phagocytosis) or 24 h (to evaluate bacterial survival) in RPMI media supplemented with 25 µg/mL gentamicin (Sigma)^41^. The phagocytic index is calculated by quantifying CFUs at 30 min and 24 h post-infection.

### Cytotoxicity Assay

Supernatants from infected cells as detailed above were used to quantify the cytosolic enzyme lactate dehydrogenase activity using the CyQuant LDH Cytotoxicity Assay (Invitrogen). The percentage of LDH activity was determined using the following formula: % of LDH release = (OD_490_-experimental LDH release OD_490_– spontaneous LDH release) / (OD_490_-maximal LDH release – OD_490_-spontaneous LDH release) x 100%^42^.

### Statistical Analysis

Data are expressed as mean ± standard error (SE) or SEM from at least three independent biological experiments performed with technical replicates. Statistical comparisons were conducted using unpaired two-tailed Student’s t-test (two groups) or one-way ANOVA with Dunnett’s multiple-comparisons test (multiple groups versus wild-type control), as specified in the figure legends. Significance was set at p < 0.05, with ns or asterisk notations denoting exact thresholds.

## ACKNOWLEDGEMENTS

This study was supported in part by grants from the NIH, including 5R21AI151571 and R01AI175461 (to J.E.V.). KT is in part supported by the Robert Austrian Research Award (RARA) by the International Society of Pneumonia and Pneumococcal Diseases (ISPPD) and Pfizer.

